# In vivo assessment of differential toxicity of cancer treatment drugs in Fanconi’s Anemia

**DOI:** 10.1101/2025.11.21.689800

**Authors:** Craig Dorrell, Sebastian Cruz-Gomez, Allison MacMillan, Alexander Peters, Barbara Burtness, Markus Grompe

## Abstract

Fanconi’s Anemia (FA) is a DNA repair disorder with a very elevated risk of cancer, especially squamous cell carcinomas (SCC). Many cancer chemotherapy agents induce DNA damage, are highly toxic in FA, and cannot be safely used in this population. The potential differential toxicity to FA patients of many new drugs being explored for use in SCC is unknown. To evaluate such compounds of unknown toxicity for use in FA cancers, we developed a sensitive in vivo bone marrow repopulation competition assay in mice. We found that afatinib, alisertib and everolimus exhibited no significant differential toxicity in this system. Therefore, these drugs are candidates for chemotherapy of cancers in human FA patients. Our competitive repopulation assay provides a robust method to screen novel chemotherapy agents for their safety in FA.

**Summary:**

‐ We developed a mouse model system for evaluating differential toxicity of cancer chemotherapy drugs on Fanconi Anemia mutant hematopoietic cells in vivo.
‐ Different drugs resulted in either increases, decreases, or neutral wildtype to
Fanconi mutant cell ratios permitting robust assessment of their safety in FA.

## Introduction

Fanconi’s Anemia (FA) is a genetic disorder characterized by impaired repair of interstrand DNA crosslinks, progressive bone marrow failure, birth defects and extreme cancer predisposition ^1,2^. Patients have mutations in any one 23 genes, encompassing the FA/BRCA pathway ^3^. The breast cancer susceptibility genes *BRCA1* ^*4*^, *BRCA2* ^5^ and *PALP* ^6^ are known members of this pathway. FA patients have a very high predisposition for acute myelogenous leukemia (AML) in childhood ^7^, but early bone marrow transplantation has significantly reduced the frequency of this lethal complication ^8^. Most FA patients now survive into adulthood, but although hematopoietic manifestations of the disease can now be treated, many adult patients are now developing solid cancers. Squamous cell carcinomas of the head and neck (HNSCC) are the most common, but many other solid tumor types also occur with a much higher frequency than in the general population ^9,10^. Unfortunately, FA cancers are very difficult to treat with the standard regimens which usually include genotoxic chemotherapy and radiation. FA patients are exquisitely sensitive to DNA damaging agents, replication stress ^11,12^ and also to inflammatory cytokines ^13-15^. Many emerging anticancer agents with promise for patients with HNSCC may have much higher toxicity in FA patients than in the general population if they increase replication stress and DNA damage either directly or secondary to immune effects. However, many new targeted agents are being developed for cancer treatment and may also be helpful for treating FA patients that have developed tumors. Although their known mechanism of action typically is not DNA damage, some of these new drugs could be differentially toxic to FA patients indirectly, for example by causing replication stress. For this reason, we developed a competitive repopulation assay in mice which provides a sensitive in vivo read-out for toxicity specific to FA cells. Wild-type and *Fancd2* mutant bone marrow cells are mixed at a 1:10 ratio and co-transplanted into a *Fancc* mutant recipients. We used the FA mutant recipient to model potential effects of drugs on the stem cell niche rather than the stem cells themselves. To establish this assay, we generated several congenic strains of mice bearing fluorescent reporter genes. This permits the use of flow cytometry to quantitate the relative contributions of stem cells from two strains of mice in our competitive repopulation assay. To date, three newer generation chemotherapy drug candidates were tested in this model. Here we found that afatinib, alisertib, and everolimus displayed no differential toxicity to FA cells and therefore may be safe to use in this patient population.

## Methods Mice

Animal care and immunization procedures were performed in accordance with the institutional review committee at Oregon Health & Science University, protocol TR03_IP00000446. 129S4 mouse strains including *Fanca*^*-/-16*^, CAGEGFP ^17^, *Fancd2*^-/-18^, and ROSA^mT/mG 19^ mutations/transgenes are maintained in our colony. CAG^EGFP^ mice were originally obtained on a c57bl/6 background and extensively backcrossed to 129S4, then bred to produce the *Fancd2*^*-/-*^/EGFP^+^ reporter strain.

### Isolation of donor cells

BM cells from CAGEGFP/*Fancd2*^-/-^ and ROSA^mT/mG^ mice were isolated from the spinal column using a streamlined version of the method of Avee Naidoo in the laboratory of Dawn Bowdish (http://www.bowdish.ca/lab/wp-content/uploads/2020/01/Isolation-of-murine-spinal-bone-marrow.pdf). Briefly, spinal columns were collected from euthanized animals and most adherent tissues removed with scissors. The clean spine was then cut into four segments and placed in a pestle containing 10 ml chilled DPBS. Four rounds of light crushing with a pestle and harvesting of liberated marrow cells through a 40 µm strainer were performed, resulting in recovery of approximately 50×10^6^ BMC per spine. Cells were aliquoted and cryopreserved in LN2 so that every experiment was performed using identical starting material.

### Competitive Repopulation

A cohort of *Fanca*^-/-^ mice were conditioned with a single dose of 600 cGy using a RadSource RS2000 X-Ray Irradiator. After 24h, each animal was briefly anaesthetized by exposure to isoflurane and retroorbitally injected with a mixture of 2×10^6^ CAGEGFP /*Fancd2*^-/-^ cells and 0.2×10^6^ ROSA^mT/mG^ (WT) cells.

### Analytical cell isolation and Flow cytometry

For pre-treatment and post-treatment assessment of peripheral engraftment, approximately 10 µl of PB was collected from the saphenous vein into tubes containing 0.1M sodium citrate. RBC were lysed in ammonium chloride prior to labeling. For post-treatment assessment of BM engraftment, cells were recovered from a femur using a standard flush protocol^20^.

Dissociated cells were resuspended at 1×10^6^ cells/ml in DMEM + 2% FBS of APC-conjugated anti-CD45 antibody (1:100, BD Pharmingen 559864) at a 1:100 dilution and incubation at 4°C for 30’. After a wash with cold DPBS, the cells were resuspended in DMEM + 2% FBS containing a DAPI (0.5 µg/ml) to label dead cells for exclusion. Cells were analyzed a BD LSRII (Becton-Dickenson, Franklin Lakes, NJ) and Flowjo X software used to quantify.

### Drug administration

Agents were administered by different routes as appropriate. MMC (RPI, Mt. Prospect IL) was delivered as a single IP injection at 0.5 mg/kg in saline, oxymetholone was provided in chow (Bioserv, Flemington NJ) formulated at 300 mg/kg (ad libitum for four weeks), and afatinib (100 mg/kg), alisertib (30 mg/kg), and everolimus (5 mg/kg) were obtained from MCE (Monmouth Junction, NJ) and administered by gavage (5 days per week for four weeks).

### Western blotting

PBMCs derived from mouse blood and cells were obtained by eliminating erythrocytes with ACK lysis buffer for 5 min at RT; cells were then washed with PBS and PBMCs were lysed using RIPA buffer for 30 min at 4 °C. Lysed cell solutions were centrifuged at 13000 rpm for 10 min at 4 °C for clarification. Supernatant was rescued and protein concentration was quantified using a BCA assay. Samples were run in an SDS-PAGE acrylamide gel at 80 V for 30 min and then 120 V for 1 h. Transference to a PVDF membrane was done overnight at 35 V, 4 °C. The PVDF membrane was blocked for 1 h at RT using 5% low-fat milk in TBST and washed three times for 5 min at RT with TBST between each step. Antibodies for western blot were incubated at 4 °C overnight and dilutions were performed using buffer 5% BSA, 0.01% azide in TBST. Antibodies used: Anti-EGFR (1:1000, CST4267), Anti-Tyr1068-phospho-EGFR (1:1000, CST3777, Cell Signaling), Anti-ERK1/2 (1:1000, CST4695), Anti-Tyr204-Tyr187-phospho-ERK1/2 (1:1000, CST5726), Anti-AURKA (1:1000, CST14475), Anti-Thr288-Thr232-Thr198-phospho-AURKA/B/C (1:1000, CST8525), Anti-CDK1 (1:1000, CST77055), Anti-phospho-CDK1 (1:1000, CST9111), Anti-Beta-Actin (1:10000, A2228, Sigma). Secondary antibody anti-rabbit-IgG-HRP (1:5000, Cytiva NA934V) or anti-mouse-IgG-HRP (1:5000, Cytiva NA931V) were incubated for 1 h at RT in 5% milk in TBST. Membrane signal was developed using WB ECL thermo-fisher kit according to manufacturer instructions using a ChemiDoc™ Imaging System from Bio-Rad.

### Statistics

The statistical significance of changes in cell type ratios pre- and post-treatment were determined by unpaired two-tailed T tests in Graphpad Prism (Boston, MA).

## Results

### Development of a competitive bone marrow repopulation assay

The hematopoietic stem cells (HSC) of FA individuals are known to be the cell type most sensitive to DNA damage and replication stress. For this reason, we decided to use competition between FA mutant and wild-type HSC during bone marrow repopulation as our toxicity assay. To account for potential effects of a FA mutant microenvironment, we used FA mutant transplant recipients (Fig.1). Three strains of mice were employed: *Fancd2* mutant mice ^18^ expressing a red fluorescent reporter, wild-type mice harboring a green reporter, and *Fancc* mutant mice ^21^ not expressing any reporter. Wild-type (green) and *Fancd2* mutant (red) bone marrow mononuclear cells were mixed at a 1:10 ratio and co-transplanted into *Fancc* mutant mice after mild conditioning (500 cGy) by irradiation. The percentage of red vs. green cells in peripheral blood was measured by FACS four weeks after transplantation to establish an engraftment baseline. As previously reported, *Fancd2* mutant cells had a significant engraftment defect, compared to wild-type controls ^22^. Despite being at a 10-fold numerical advantage in the transplant donor cells, the ratio of wild-type/*Fancd2* blood was ∼ 1:1 after 4 weeks.

**Figure 1.**
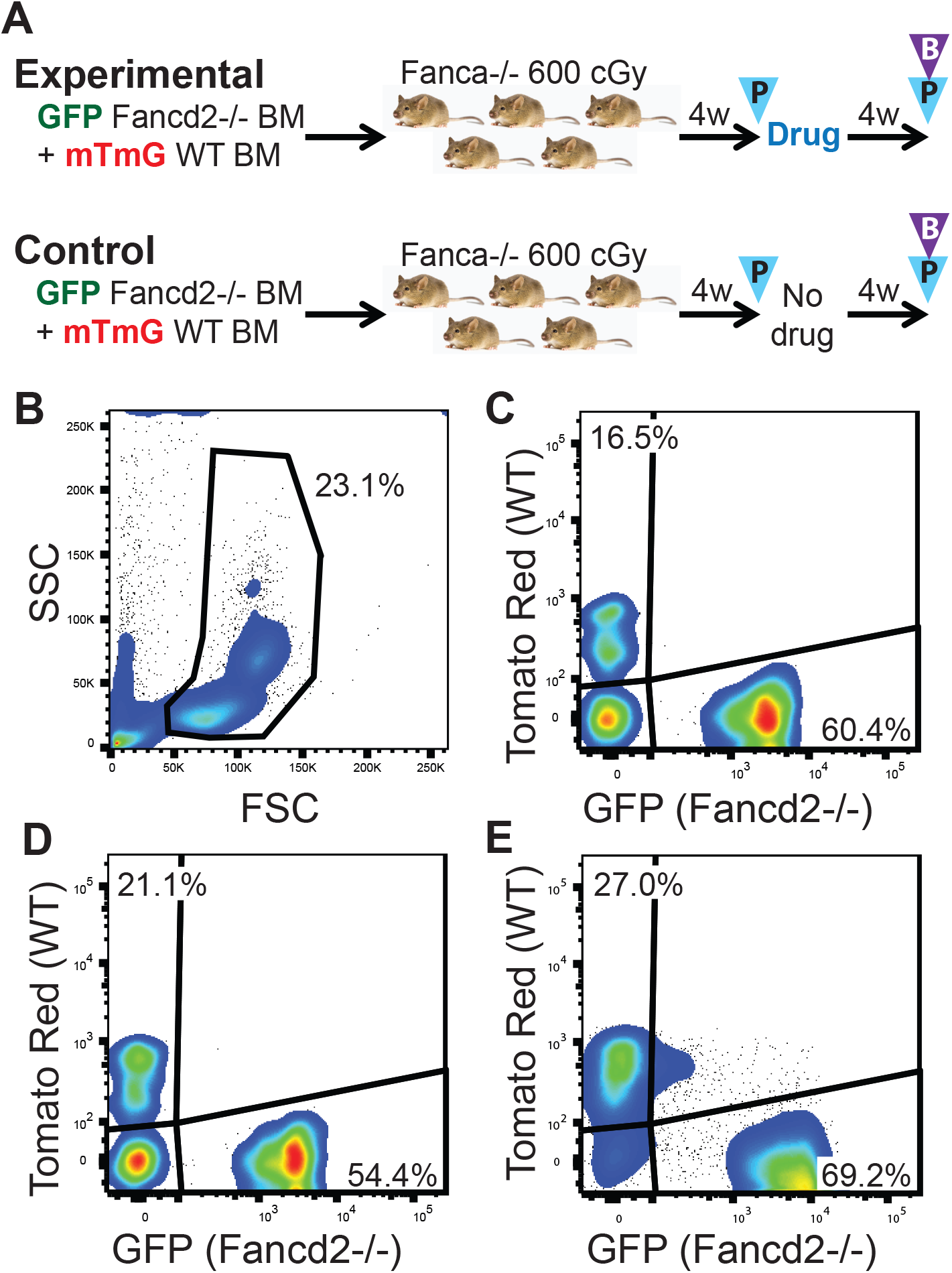
Experimental scheme. (A) Competitive repopulation was monitored in mouse cohorts before and after drug administration by sampling peripheral blood (blue triangles) and bone marrow (purple triangle). After FSC/SSC (B) and CD45+/DAPI-gating, the frequencies of mutant (green fluorescent) and WT (red fluorescent) were measured before (C) and after (D) treatment in peripheral blood. Bone marrow (E) was assessed at terminal harvest to assess progenitor survival. C,D and E are from a no-treatment control and showed no significant shift in wt/mutant cell ratios over time.

Next, half the animals were treated with the test agent for 4 weeks and half served as non-treatment controls. Controls and experimental transplant recipients were age- and sex matched. After completion of the drug treatment, peripheral blood was re-analyzed. After an additional 4 weeks without drug treatment, terminal harvest and analysis was performed. The ratio of wt/FA cells in peripheral blood and bone-marrow was determined by FACS (Fig.1).

### Effects of specific drugs

To validate our ability to detect FA-specific toxicity in this assay, we used mitomycin C as a control. MMC is a potent DNA cross-linker and known to be very toxic to FA cells and individuals ^23-25^. As expected, treatment with MMC dramatically reduced the contribution of *Fancd2* mutant stem cells to repopulation by over 100-fold, thus confirming the ability of system to detect toxicity against FA stem cells (Fig. 2). Next, we explored the effects of 3 drugs with potential to be used in cancer therapy of FA patients ^26,27^. All of them have been used in non-FA cancer patients all have a primary mechanism of action distinct from DNA damage, although effects on replication stress and inflammation may be incompletely characterized. We tested afatinib ^26-28^, an EGFR-family antagonist, alisertib ^29,30^, an Aurora kinase A inhibitor, and everolimus ^31,32^, a blocker of mTOR signaling. We used previously published doses of the compounds ^33-35^, thought to be fully effective in mice. To confirm the in vivo activity of these drugs in FA mice, we performed Western blots of the target genes of afatinib and alisertib (Fig.3). As expected, these assays confirmed the biological activity of the compounds.

**Figure 2.**
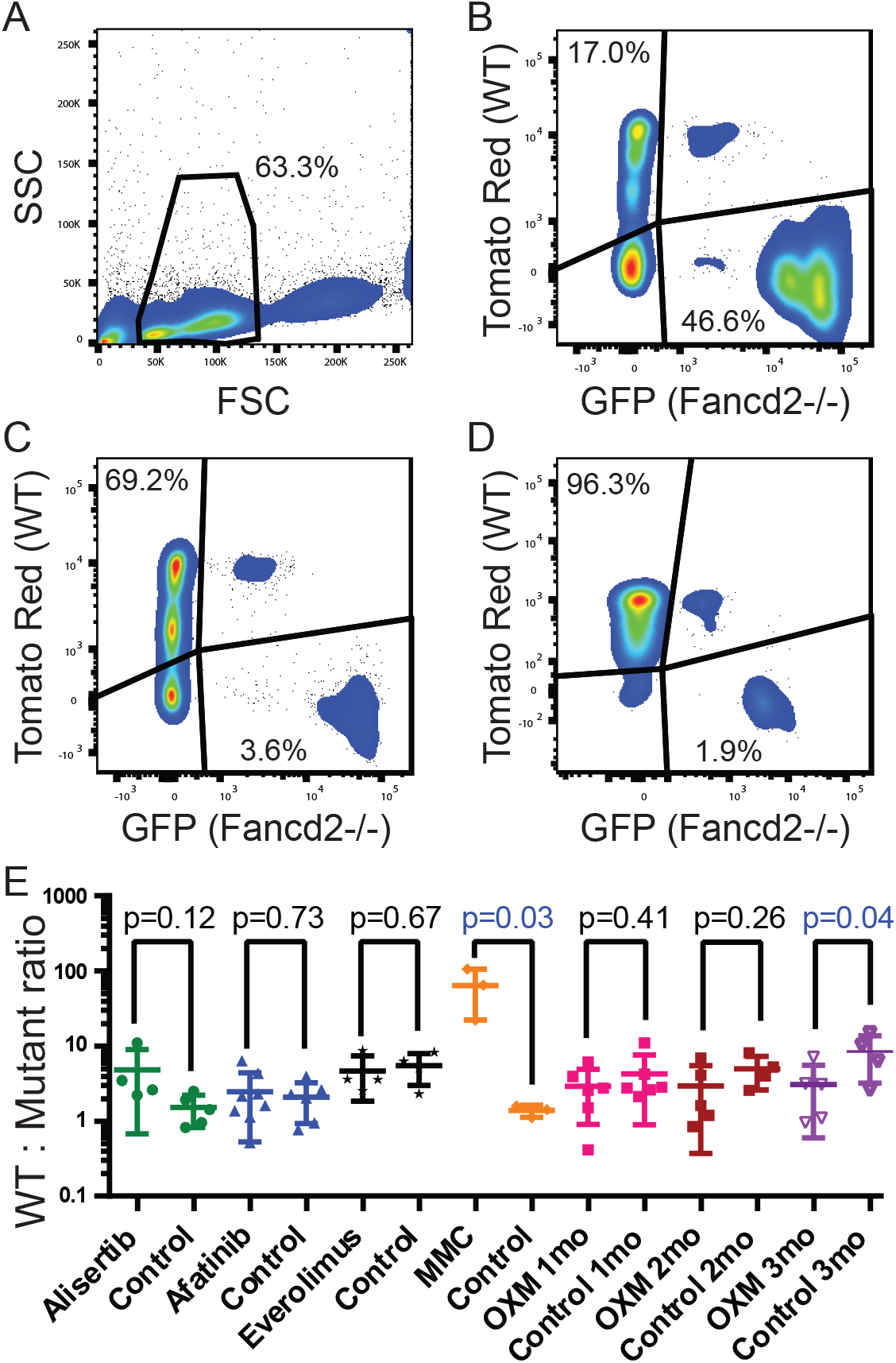
Drug treatment can dramatically alter cell type ratios during competitive repopulation. After FSC/SSC (A) and CD45+/DAPI-gating, a comparison of pre-(B) and post-(C) MMC treatment shows a dramatic reduction in the contribution of mutant cells to peripheral blood. BM contained very few mutant cells at harvest (D). Changes in the ratios of WT to *Fancd2*^-/-^ cells in peripheral blood after 4 weeks of drug treatment (E). For each treated or control animal, values shown are the ratio of the ratio between the cell types pre-treatment (1 month after engraftment) and post-treatment (2 months after engraftment) as measured by flow cytometric analysis of blood.

**Figure 3.**
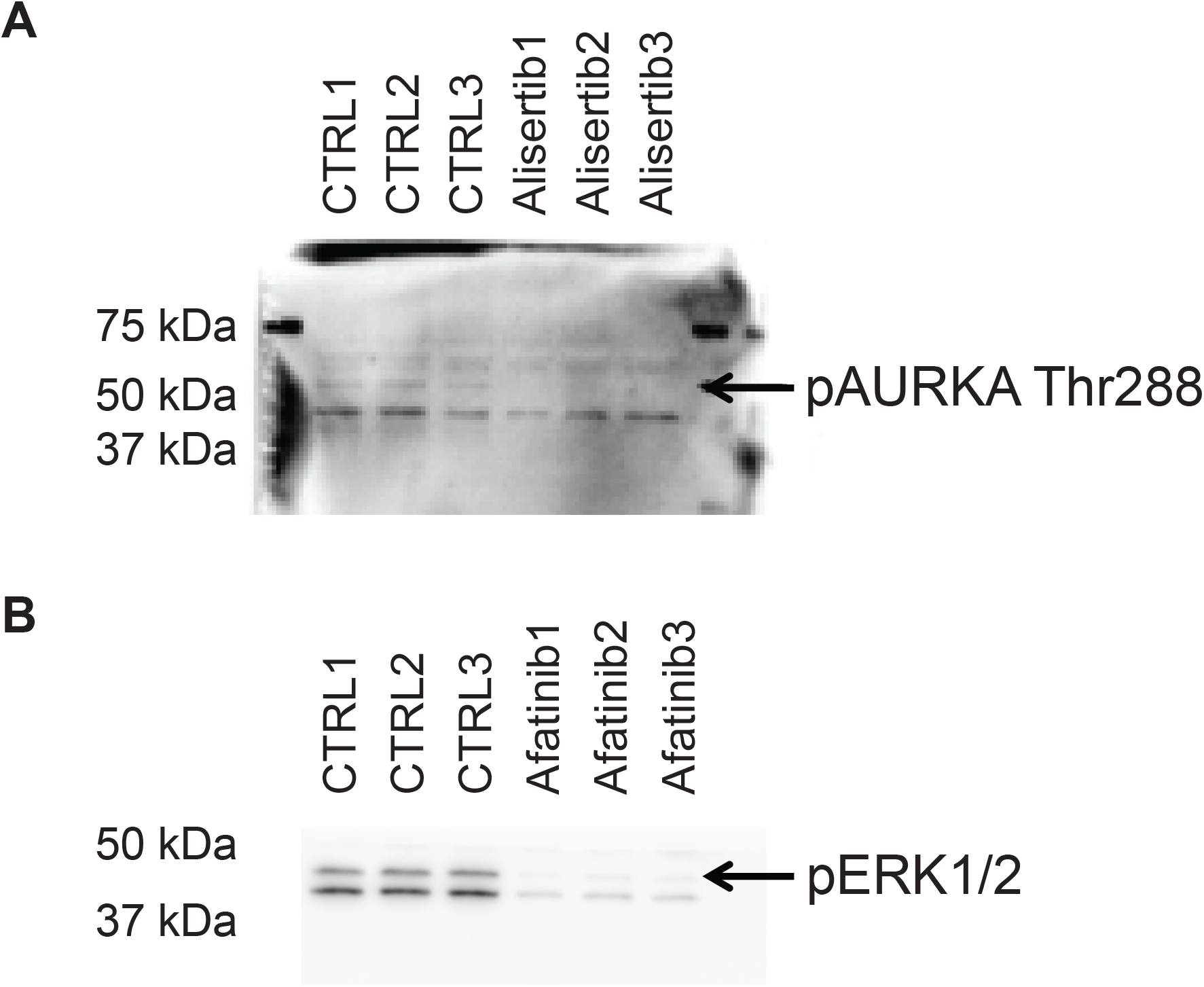
Western blotting of protein lysates from peripheral blood collected at final harvest. Membranes were labeled with anti-phosphoAURKA thr286 to measure downregulation of Aurora kinase by Alisertib (**A**) or anti-phosphoERK1/2 (**B**) for downregulation of EGFR by Afatinib. The relevant protein bands are indicated with an arrow.

Gratifyingly, none of the 3 candidate drugs had a significant differential effect on the function of FA HSC, suggesting that each would be safe to use in the clinical setting.

The ratio of Fanconi to wild-type blood cells was similar in drug-treated animals and placebo controls. The results obtained from peripheral blood analysis were similar to those obtained by analyzing bone marrow (Fig.2).

As a further validation of our assay, we also studied the effect of oxymetholone, a drug that is used to treat bone marrow failure in FA patients ^20,22,36^. Anabolic steroids are known to increase blood counts in FA patients ^37^. Interestingly, the drug enhanced the ratio of *Fancd2* to wild-type engraftment, consistent with a differentially positive effect on FA stem cells vs. wild-type controls (Fig.2). This finding indicates that our assay is also suitable to detect drugs that are beneficial to FA cells, improving their engraftment, survival, or expansion.

## Discussion

Solid cancers are very difficult to treat with conventional chemotherapy in Fanconi Anemia patients. Many case reports of death from standard doses of anti-tumor agents in FA exist ^38,39^. The typical conditioning regimen for bone marrow transplantation is also too toxic in this population ^40-42^. These observations have created the imperative to avoid genotoxic agents and treat patients with only surgery or chemotherapy drugs that are not thought to be genotoxic. In bone marrow transplantation, antibody mediated conditioning has recently been shown to be an alternative to total body irradiation in FA ^43^. Many of the newer agents for solid cancer treatment are aimed at targets that are apparently unrelated to DNA damage and repair. These compounds are excellent candidates to be used in FA patients. However, FA cells are not only susceptible to outright DNA damage, but also to inflammatory cytokines and stalling of replication forks ^11,12^. Given the irreversible and potentially catastrophic effects of overdoses in this unique population, our in vivo assay can derisk the application of drugs that have never been used in FA patients. We were not surprised that the EGFR family inhibitor afatinib was not toxic, but we did consider the possibility that the aurora kinase inhibitor alisertib might cause replication stalling and be problematic. Gratifyingly, this was not the case. In addition, everolimus also appeared safe.

Our positive control for toxicity, mitomycin C, gave the expected result, as did our control for therapeutic benefit, oxymetholone. We therefore propose that our assay should be used before a new non-genotoxic agent is employed in human FA patients.

It is worthwhile pointing out that we used fully immune competent animals in our test system. Therefore, any indirect DNA damage or replication stress caused by immunological effects of a candidate drug should be captured by this assay.

Our positive result with an anabolic steroid, known to improve stem cell performance in FA, highlights the utility for our assay in assessing the effects of small molecules with potential beneficial effects in FA.

## Acknowledgements

This work was supported by grants from the Fanconi Anemia Cancer Fund (https://fanconi.org) and Stand-up to Cancer Foundation (https://standuptocancer.org). The OHSU Flow Cytometry and Monoclonal Antibody Shared Resource (RRID:SCR_009974) provided instruments for essential cell analysis and the OHSU department of Comparative Medicine for helping to maintain our mouse colony.

## Authorship

Contribution: M.G., B.B., and C.D. designed the research; C.D., S.C.-G., A.M., and A.P. performed the research; C.D. and S.C.-G. collected the data; C.D. and S.C.-G. analyzed and interpreted the data; C.D. performed statistical analysis; and M.G., C.D., S.C.-G., and B.B. wrote the manuscript.

